# Variation in repeat copy number of the *Epithelial adhesin 1* tandem repeat region leads to variable protein display through multiple mechanisms

**DOI:** 10.1101/872853

**Authors:** Colin J. Raposo, Kyle A. McElroy, Stephen M. Fuchs

**Affiliations:** Department of Biology, Tufts University, Medford, MA 02155; Allen Discovery Center, Tufts University, Medford, MA 02155

**Author notes:** Corresponding Author: 200 Boston Ave Suite 4700 Medford, MA 02155.

## Abstract

The pathogenic yeast *Candida glabrata* is reliant on a suite of cell surface adhesins that play a variety of roles necessary for transmission, establishment, and proliferation during infection. One particular adhesin, Epithelial Adhesin 1 [Epa1p], is responsible for binding to host tissue, a process which is essential for fungal propagation. Epa1p structure consists of three domains: an N-terminal intercellular binding domain responsible for epithelial cell binding, a C-terminal GPI anchor for cell wall linkage, and a serine / threonine-rich linker domain connecting these terminal domains. The linker domain contains a 40-amino acid tandem repeat region, which we have found to be variable in repeat copy number between isolates from clinical sources. We hypothesized that natural variation in Epa1p repeat copy may modulate protein function. To test this, we recombinantly expressed Epa1p with various repeat copy numbers in *S. cerevisiae* to determine how differences in repeat copy number affect Epa1p expression, surface display, and binding to human epithelial cells. Our data suggest that repeat copy number variation has pleiotropic effects, influencing gene expression, protein surface display, shedding from the cell surface, and host tissue adhesion of the Epa1p adhesin. Understanding these links between repeat copy number variants and mechanisms of infection provide new understanding of the variety of roles of repetitive proteins contribute to pathogenicity of *C. glabrata*.

## Introduction

Systemic infections by yeast of the *Candida* genus, or invasive candidiasis, have been an emerging public health concern since the 1980s and are a particular concern to hospitalized patients suffering from immune deficiencies such as HIV/AIDS (Fidel *et al.*, 1999, Lin *et al.*, 2001, McNeil *et al.*, 2001, Pfaller & Diekema, 2007). *Candida glabrata* follows *Candida albicans* as the second most common cause of invasive candidiasis in the United States and tends to cause particularly severe infections as a result of ever-increasing rates of antifungal resistance (Pfaller & Diekema, 2007, Alexander *et al.*, 2013, Spettel *et al.*, 2019). In addition, *C. glabrata* infections are made worse by the highly plastic haploid genome that allows for the pathogen to adapt to a wide range of environments, such as those experienced in different tissues and on various abiotic surfaces (Fidel *et al.*, 1999, Ahmad *et al.*, 2014, Lopez-Fuentes *et al.*, 2018).

There are over 60 cell-wall associated proteins, members of a large radiation of the adhesin superfamily, many of which are essential for the establishment and propagation of *C. glabrata* infection (Timmermans *et al.*, 2018). These adhesins can bind a variety of substrates and serve many functions including transmission of the pathogen, establishment of infection, and formation of antibiotic-resistant biofilms. First identified by Cormack and colleagues, deletion of the gene which codes for one prominent adhesin, Epithelial Adhesin 1 [*EPA1*, Epa1p], results in a 95% reduction in adherence to cultured epithelial cells. Heterologous expression of *EPA1* in *Saccharomyces cerevisiae* confers the ability of cells to bind to epithelium (Cormack *et al.*, 1999). Epa1p-mediated binding to host epithelial cells is reliant on the interaction between the N-terminal Epa1p lectin-like binding domain and a terminal galactose β1-3 glucose residue of a host cell surface glycan (Maestre-Reyna *et al.*, 2012).

To facilitate binding, Epa1p structure enables the presentation of the lectin-like domain (A domain) on the cell surface. A hydrophobic N-terminal signal peptide is located at amino acid residues 6-24 and is responsible for the translocation of the protein into the secretory pathway and eventual targeting to the cell surface (Schekman, 1982, Cormack *et al.*, 1999). Once at the surface, Epa1p is anchored by a C-terminal glycosylphosphatidylinositol (GPI) anchor, which is responsible for covalent linkage of the protein to cell wall glycans (Cormack *et al.*, 1999, Frieman *et al.*, 2002, Klis *et al.*, 2010).

Epa1p contains a serine/threonine-rich linker (B domain) of roughly 600-amino acids between the binding domain and GPI anchor (Cormack *et al.*, 1999). This domain has been shown to be highly modified by both O- and N-linked glycosylation (Frieman *et al.*, 2002). This is a process which is predicted to further crosslink GPI-linker cell wall proteins (GPI-CWP) to the glycan network of the cell wall to affect protein structure and function (Frieman *et al.*, 2002, Klis *et al.*, 2010). Based on models of other GPI-CWPs, glycosylation of Epa1p has been predicted to stabilize the protein into an extended rod structure to enhance A-domain clearance of the cell wall (Jentoft, 1990, Frieman *et al.*, 2002). Alternatively, stability of Epa1p in the cell wall may be regulated in a glycosylated-mediated mechanism similar to Flocculation 11 [Flo11p], a related GPI-CWP in *S. cerevisiae*. When Flo11p is under-glycosylated, its stability in the cell wall decreased compared to controls as a result of increased susceptibility to extracellular proteases such as Kexin 2 (Karunanithi *et al.*, 2010, Meem & Cullen, 2012).

Within the ser/thr-rich B domain of Epa1p, there is a tandem repeat region of a 40-amino acid sequence (T/M)VRSTLPSSAGSNETSINVPFSSSTESNTSTSSTSTSNSK, which repeats three times in the reference genome of *C. glabrata* (Cormack *et al.*, 1999). DNA minisatellite sequences, such as that which codes for this repetitive domain, are known to be genetically unstable and prone to expansions and contractions (Richard & Dujon, 2006, Gemayel *et al.*, 2010, Thierry *et al.*, 2010). Variation in repeat regions, both in repeat copy number and mutations within repeats, has been shown to have an important impact on phenotype and serve as loci for population-wide genetic variability (Babokhov *et al.*, 2018, Babokhov *et al.*, 2018). While six of the nine genes in the EPA family contain repetitive regions, the link between repeat variation and function of these proteins has not been widely characterized (Thierry *et al.*, 2010, Ahmad *et al.*, 2014). However, increased repeat copy number in the *FLO* genes, which code for similar GPI-CWPs in *S. cerevisiae*, improves adherence, so it has been postulated that a similar effect will be seen in the Epa adhesins (Verstrepen *et al.*, 2005, Fidalgo *et al.*, 2006, Thierry *et al.*, 2010). While studies of Epa1p have not yet specifically focused on the repeat region, the overall size of the linker domain is positively correlated to Epa1p-mediated binding to epithelial cells (Frieman *et al.*, 2002). As discussed above, there may be a variety of mechanisms at play resulting in this enhanced adhesion conferred by adhesin proteins with extended linker domains including increased clearance of the cell wall and higher protease protection of longer repeat variants.

In this study, we aimed to identify whether repeat copy number variation exists in the *EPA1* gene across *C. glabrata* isolates collected from a variety of clinical samples and determine whether this variation influenced Epa1p function. Here, we show that several length variants exist, and that in a clinical population, repeat variants of between three and five repeats are most common. We determine that repeat copy number variation has complex effects on protein function. In our recombinant system, the effect of repeat copy number on the surface display of Epa1p occurs through modulation of both the expression of *EPA1* and stability of Epa1p at the cell surface. In sum, these data show a link between repeat copy number variation and *C. glabrata* pathogenicity, which potentially offers a new angle for therapeutic intervention during *C. glabrata* infections.

## Methods

### Yeast Strains and Clinical Isolates

Clinical isolates of *C. glabrata* were provided by Dr. Yoav Golan from the Tufts Medical School (Boston, MA). Strains were named numerically (CG1-CG24; Table S1). The *C. glabrata* reference strain BG14 (ura3::Tn903 G418^R^; (Cormack & Falkow, 1999)) was a gift from Professor Brendan Cormack (Johns Hopkins University). For all experiments involving *S. cerevisiae*, strain BY4741 (MATa his3Δ1 leu2Δ0 met15Δ0 ura3Δ0) was used. BY4741 and BG14 were grown at 30°C in YPD media supplemented with 2% agar when required. For experiments requiring *S. cerevisiae* carrying plasmid DNA, strains were grown with Synthetic Complete dropout media lacking one or more amino acids for selection, supplemented with 2% agar when required. For short term usage, cultures were kept on plates at 4 °C. For long term storage, cultures were supplemented with 15% glycerol, and cells were stored at −80 °C.

### HeLa Cell Culture

Human cervix adenocarcinoma (HeLa) cells were cultured in Dulbecco’s Modified Eagle Medium supplemented with 10% heat inactivated fetal bovine serum. Cells were cultured at 37°C with 5% CO_2_ and were passaged every 2–4 days before reaching 100% confluence.

### Analysis of EPA1 repeat variation

Genomic DNA was isolated from *C. glabrata* strain through a rapid yeast genomic prep (Hoffman & Winston, 1987). We then amplified the repetitive region of *EPA1* through PCR with primers EPA1repF and EPA1repR (see Table S2 for primer sequences) which anneal to *EPA1* 36 and 58 bp upstream and downstream of the repeat region, respectively. The DNA isolated from the repeat region was then analyzed by gel electrophoresis to determine repeat copy number and confirmed by sequencing. For DNA sequencing analysis, we isolated the *EPA1* gene from *C. glabrata* genomic DNA with the OCR014 and OCR015 primers, and the sequencing was carried out by Eton Biosciences Inc. (Charlestown, MA) with the EPA1repR primer. Sequences were then aligned utilizing the translation alignment tool in Geneious Prime (Biomatters Ltd).

### Plasmid construction

For expression of *EPA1* in *S. cerevisiae*, the pSPHA expression vector was used. (See Figure S1 for plasmid maps and cloning strategy.) The pSPHA expression vector allows for constitutive expression of a protein product with an N-terminal signal peptide from Epa1p and 3x hemagglutinin (HA) tag. The vector was maintained in *S. cerevisiae* by growth in Synthetic Complete minus Leucine media and an ampicillin resistance marker allows for selection and cloning in *Escherichia coli*. To build the pSPHA expression vector, the pRS415 expression vector (pADH; Addgene 87374) was modified through the addition of the 254 bp SPHA cassette (GeneArt; Table S2) coding for the signal peptide, 3x HA tag, and a 3’ multiple cloning site into the SmaI (EC 3.1.21.4) restriction site via Gibson Assembly. The final construct was confirmed by DNA sequencing.

To build the *EPA1* expressing construct pSPHA-EPA1, the fragments of the *EPA1* gene containing the A, B, and GPI domains were isolated using the OCR035 and OCR036 primer set. Four different *EPA1* fragments were created; *EPA1* was amplified from BG14, CG3, CG15, and CG2 with three, four, five, and ten repeats respectively [*EPA1* (3 rep), *EPA1* (4 rep), *EPA1* (5 rep), *EPA1* (10 rep); Figure 1C]. These fragments were then inserted into SmaI-linearized pSPHA through Gibson Assembly to create the final expression vector, which was confirmed via sequencing. pSPHA-EPA1(0 rep) was created by deleting the repeat region from *EPA1* by QuickChange Mutagenesis (Agilent), amplifying *EPA1* with OCR035 and OCR036 primers and cloning via Gibson Assembly as described above.

**Figure 1.**
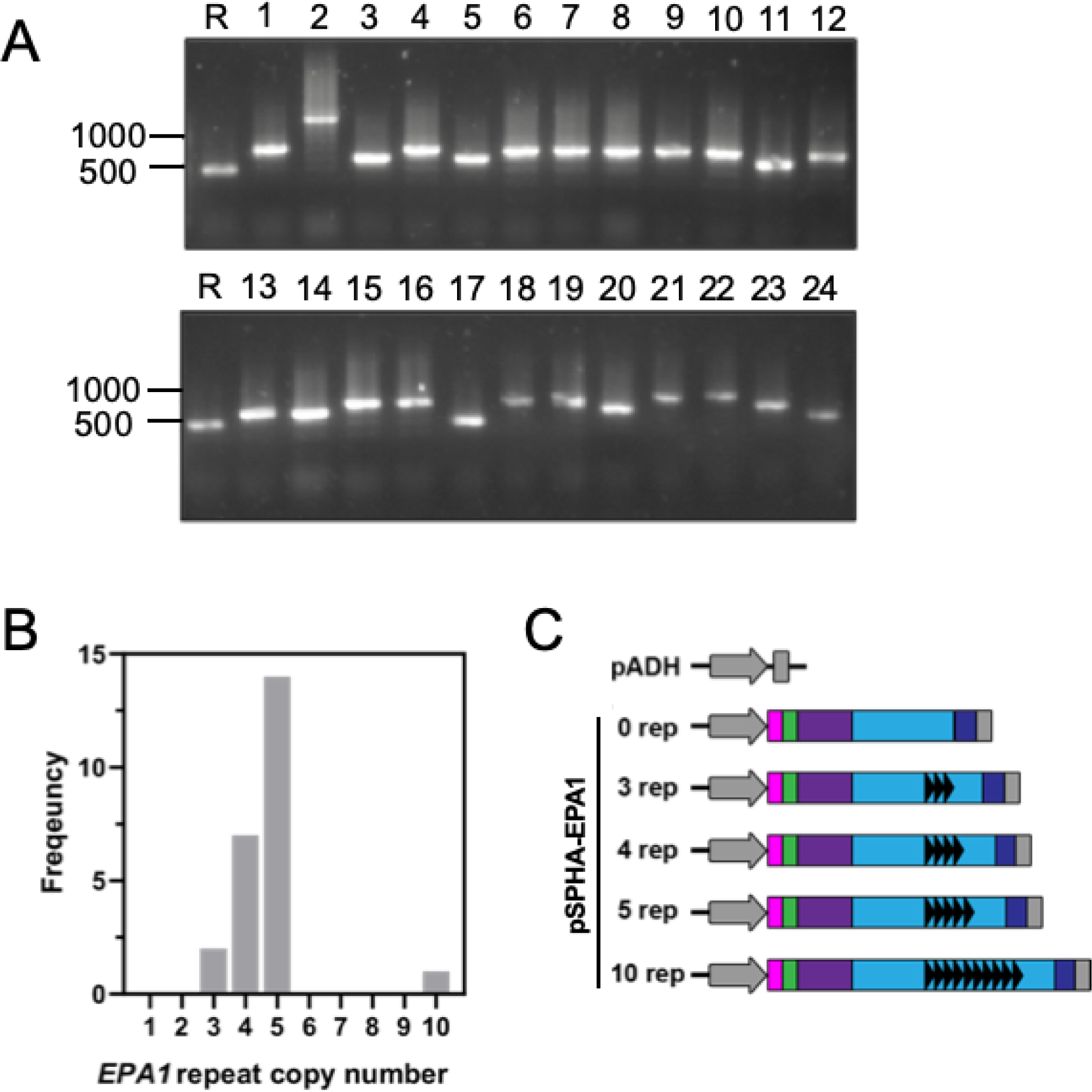
EPA1 codes for an adhesin with variable repeat copy number. **(A)** *EPA1* repeat region amplified from gDNA of 24 *C. glabrata* clinical isolates (1-24) and BG14 reference strain (R) through PCR and visualized on an agarose gel. **(B)** *EPA1* repeat copy number frequency from the 24 clinical isolates. **(C)** pSPHA-EPA1 expression vector cloned into *S. cerevisiae* allows for constitutive expression of HA-tagged Eap1p (A domain - purple, B domain – light blue, GPI anchor – dark blue) with a 3x HA-tag (green) directed to the surface by the Epa1p signal sequence (pink) under the control of the ADH promoter (grey arrow) and CYC1 terminator (grey box). Five repeat copy number variants of *EPA1* were cloned; the black arrows represent each individual tandem repeat. pADH is the empty vector.

### Flow Cytometry Analysis

Measurement of surface display by HA-tagged Epa1p was achieved through extracellular staining of cells and flow cytometry analysis. pSPHA-EPA1 carrying strains were grown to log-phase and stained with chicken anti-HA antibody (Gallus Immunotech AHA) in PBS + 0.1% BSA (PBSA). Cells were then stained with AlexaFluor 488 conjugated goat anti-chicken secondary antibody (Abcam 150169) in PBSA. Stained cells were resuspended in PBSA and analyzed on an Attune NxT Flow Cytometer (Thermo Fisher).

The flow cytometry gating strategy is outlined in Figure 2A. Briefly, debris and doublets were gated out, and 10,000 events within the singlet population were randomly selected through the FlowJo DownSample plugin (FlowJo, LLC). The BL-1 laser (excitation 488nm/emission 530nm bp 30nm) was used to measure AlexaFluor 488 signal, and the population of HA+ cells was determined by setting the gate on the pADH negative control. From this population we determined the percentage of singlets stained for HA (%HA+) and the mean fluorescent intensity of the HA+ population (MFI).

**Figure 2.**
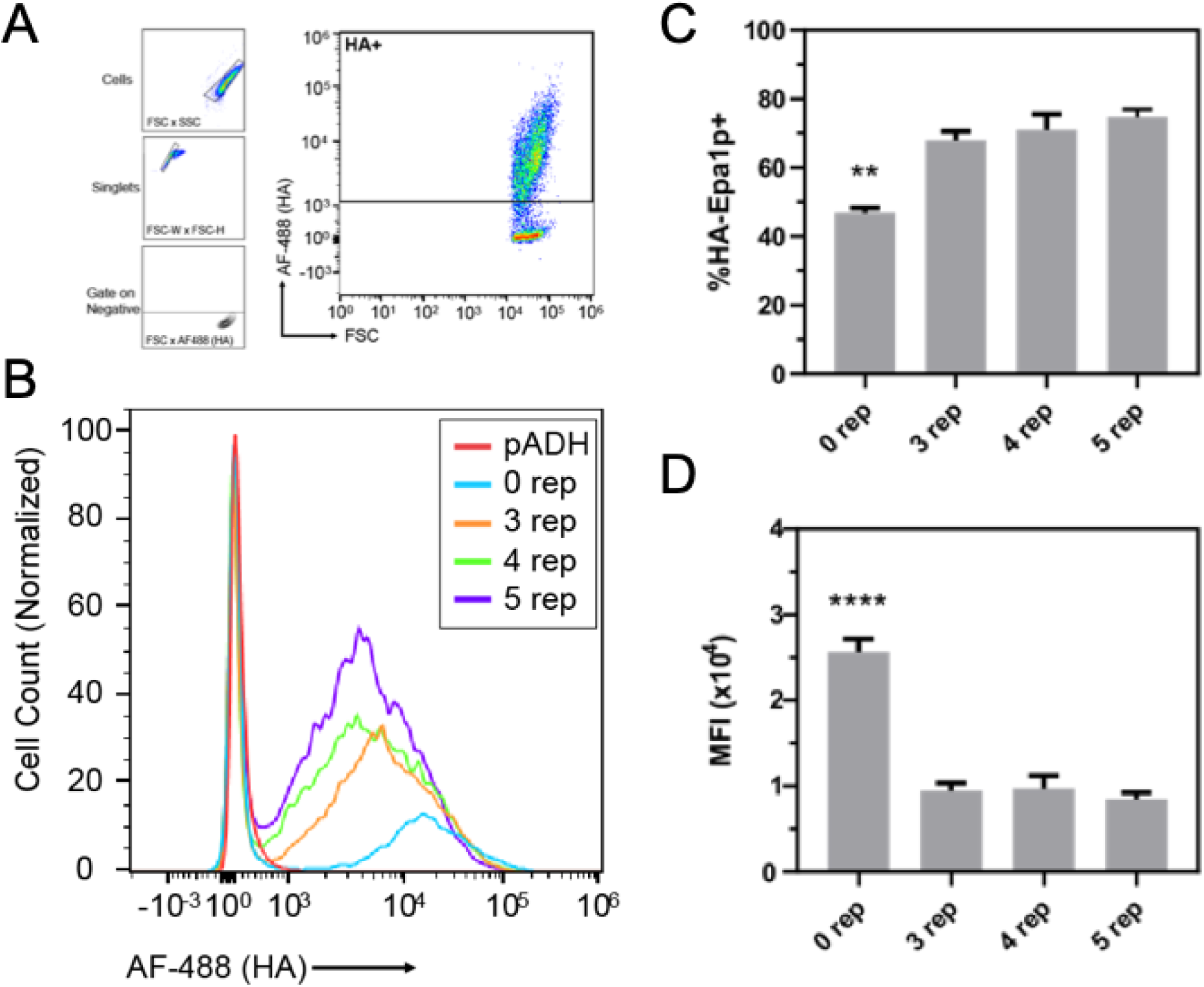
Epa1p repeat copy number affects surface display. HA-Epa1p with 0, 3, 4, and 5 repeats was expressed in *S. cerevisiae* under constitutive expression under the ADH promoter, and surface display was measured via □HA flow cytometry. **(A)** Flow cytometry gating strategy. Single cells were selected from debris and doublets, and the negative control (pADH) was utilized to set the gate for the HA+ population detected by positive AlexaFluor 488 fluorescence. **(B)** Surface display of HA-Epa1p with various repeat copy numbers. Y-axis counts are normalized by division by the count of the mode fluorescent intensity of each sample. Plot is representative of multiple experiments. **(C)** Percentage of cells stained positively for HA-Epa1p. **(D)** Mean fluorescence intensity (MFI) of HA+ stained cells. Mean + SEM is shown. n=3. Representative experiment from multiple independent experiments. ** = p<0.01, **** = p<0.0001 as determined by paired Bonferroni corrected T-tests between all experimental groups.

**Figure 3.**
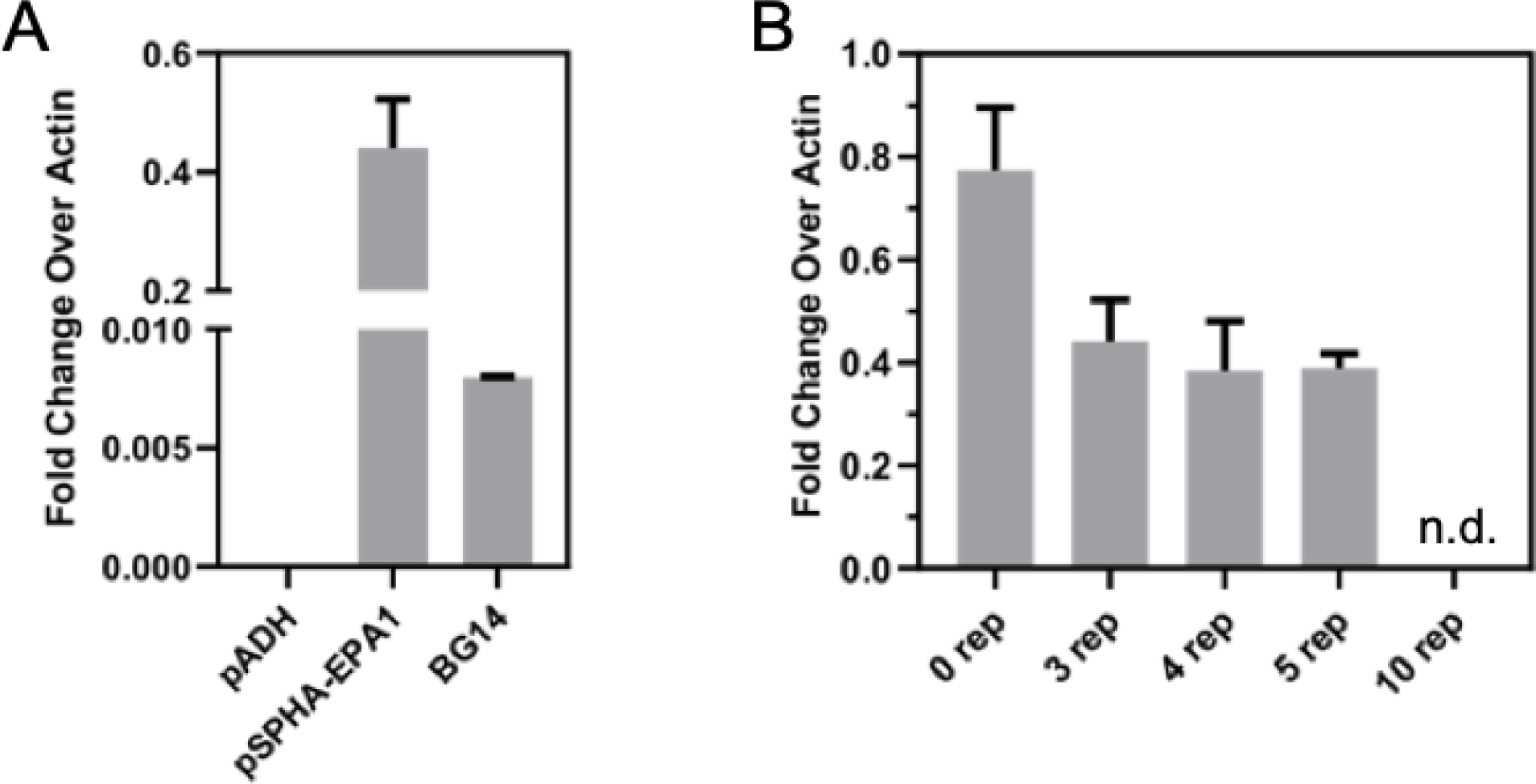
EPA1 mRNA expression levels decrease with increased repeat copy number. **(A)** mRNA expression levels of *EPA1* in *S. cerevisiae* carrying the empty pADH vector or pSPHA-*EPA1* (3 rep) or BG14, a lab strain of *C. glabrata* measured by rt-qPCR. Expression is expressed as fold change over Actin (*ACTI*). **(B)** RT-qPCR-quantified expression levels of *HA-EPA1* with various repeat copy numbers (0, 3, 4, 5, 10) of the *EPA1* repeat region expressed through the pSPHA-*EPA1* plasmid system. n.d.= no detection. Expression is expressed as fold-change over expression of Actin (3 rep). Mean + SEM is shown. n=2. Representative experiment from multiple independent experiments.

### Reverse Transcriptase Quantitative PCR

To measure levels of *EPA1* expression at the mRNA level, yeast were grown to log phase and spheroplasted in 0.02 U/mL zymolyase in 1M sorbitol + 100mM EDTA; total RNA was extracted with the illustra RNAspin Mini Kit (GE Healthcare). Complementary DNA was then synthesized using the SuperScript III First Stand Synthesis System (Invitrogen) with random hexamer primers. Quantitative PCR (qPCR) was then performed with SYBR Green 2x Master Mix (Applied Biosciences) or Brilliant III Ultra-Fast SYBR Green QPCR Master Mix (Agilent) utilizing the EPA1qPCR F and EPA1qPCR R primer set to amplify the *EPA1* amplicon, and the cycle threshold (C_T_) was determined within the linear range of amplification. C_T_ values for each reaction were then normalized to internal *ACT1* controls, amplified with ACT1qPCR F and ACT1qPCR R primers.

### Detection of Shed Protein via Western Blot

To analyze the shedding profile of HA-Epa1p from the yeast cell wall, we collected and analyzed growth media through Western Blotting (Figure 4A). To prepare samples, pSPHA-EPA1 carrying strains were grown to log phase and equivalent volumes of culture based on cell OD_600_ were collected. Cultures were then spun down; the media supernatant was collected (cells were discarded) and immediately frozen at −80 °C. Frozen media samples were lyophilized to a film and reconstituted at 40x concentration in 50mM Tris, 5mM EDTA, pH8 buffer. Laemmli SDS-PAGE sample buffer was added, and the sample was heated for 5 minutes at 95°C.

**Figure 4.**
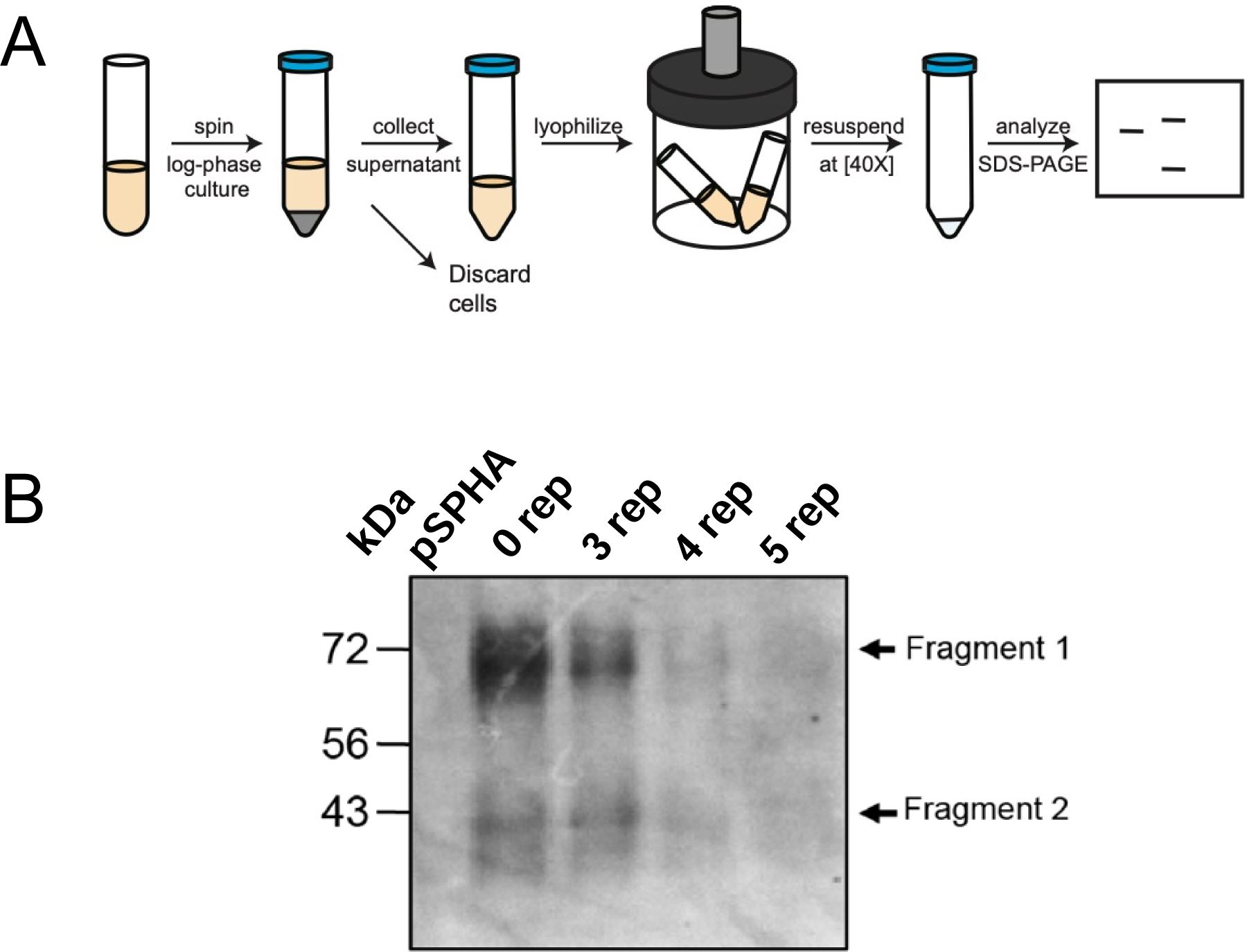
Shedding of HA-Epa1p fragments increases with reduced repeat copy number. **(A)** HA-Epa1p shedding assay. Briefly, cells were grown to log phase and their growth media was collected by centrifugation. Media was then concentrated by lyophilization and resuspended at 40x in Tris-EDTA. **(B)** □HA Western Blot of concentrated media from cells expressing HA-Epa1p with various repeat copy numbers (0, 3, 4, 5 rep) or the empty pSPHA vector. Media were collected from an equivalent number of cells and concentrated via lyophilization before Western Blot analysis. Blot is representative of three replicates.

The supernatant from the media collected from 9×10^6^ cells was run on a 10% SDS-PAGE gel and transferred to a PVDF membrane (GE Lifesciences). The membrane was probed with monoclonal mouse anti-HA antibody (Invitrogen 326700) freshly diluted (1:333) in PBST + 2.5% non-fat milk followed by incubation with HRP-conjugated goat anti-mouse secondary antibody (Invitrogen 626520) in PBST + 2.5% non-fat milk (1:1000). Signal on the blot was detected with Amersham ECL Prime Western Blotting Detection Reagent (GE Lifesciences) and imaged with autoradiography film (Andwin Scientific).

### Epithelial Cell Adherence Assay

The adherence of various yeast strains to epithelial cells was measured through a binding assay based on those outlined by (Cormack *et al.*, 1999, Halliwell *et al.*, 2012, Diderrich *et al.*, 2015; Figure 5A). First, HeLa cells were grown to confluence in a 24-well dish and fixed with 2% formaldehyde in phosphate buffered saline (PBS). After fixation, the HeLa cells were kept in HeLa culture media treated overnight with 0.2 U/mL of □2-3,6,8,9 Neuraminidase (New England Biolabs) to cleave sialic acid from the cell surface glycans.

**Figure 5.**
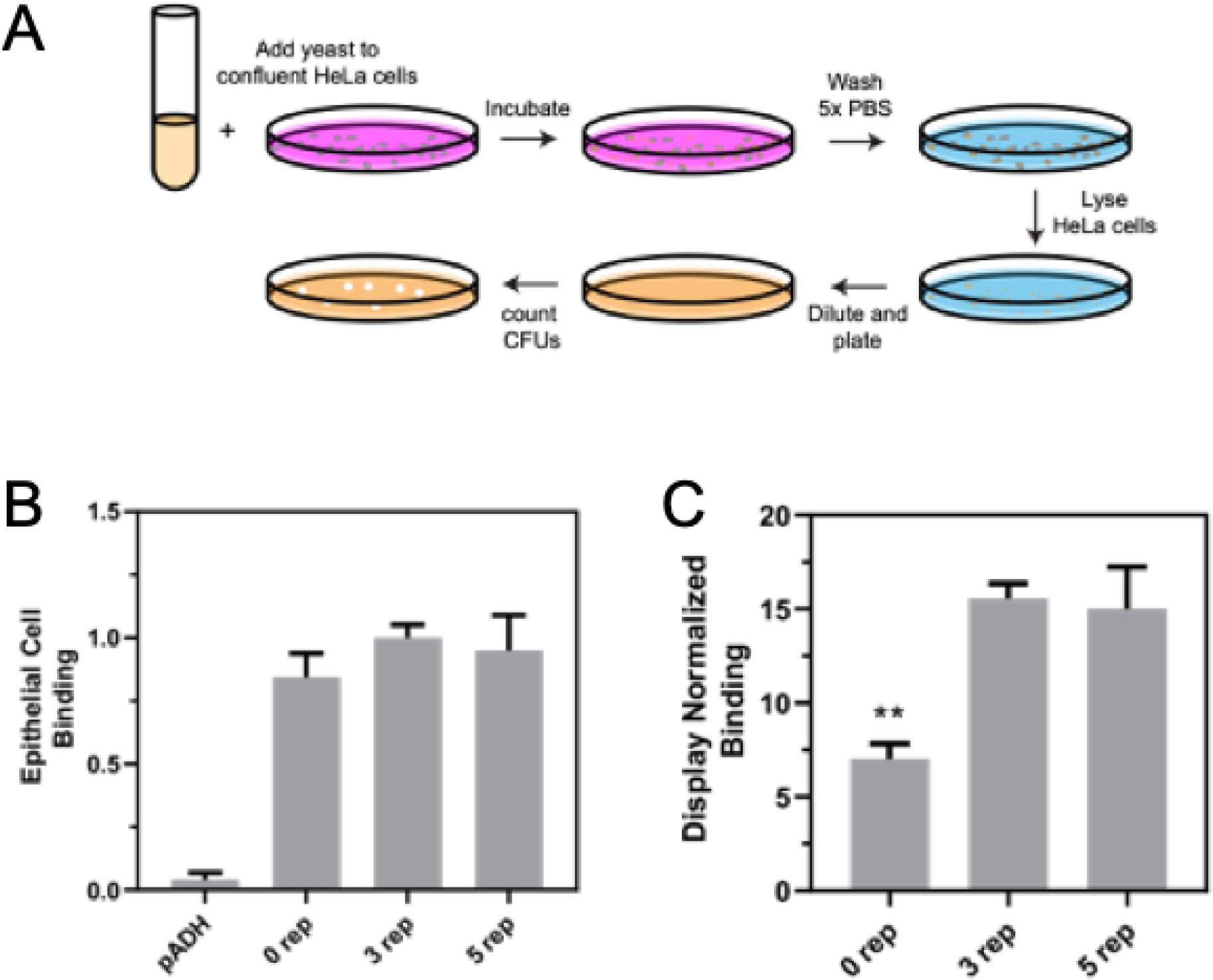
Quantifying HA-Epa1p mediated epithelial cell adherence. **(A)** Binding of *S. cerevisiae* expressing HA-Epa1p repeat copy number variants (0, 3, 5 repeats) to epithelial cells was measured via an epithelial cell adherence assay. Unbound yeast were then removed via washing in PBS, and HeLa cells were chemically lysed. Bound yeast were quantified by counting colony forming units (CFUs) and comparing them to the CFUs on input control plates. **(B)** Binding normalized to HA-Epa1p (3 rep). **(C)** Display normalized binding, which normalizes binding to surface display metrics determined via flow cytometry (percent positive and Mean Fluorescence). Values are mean + standard deviation. n=5 for 5 rep, n=6 for others. Pooled from multiple experiments. ** = p<0.01 as determined by paired Bonferroni corrected T-tests between all groups.

To adhere yeast to HeLa cells, *S. cerevisiae* carrying pSPHA-EPA1 with various repeat copy numbers or carrying the pADH empty vector were grown to log phase and 1×10^6^ yeast cells were added to treated HeLa cells in fresh HeLa culture media. The co-culture was spun for 2 min at 200 rcf to induce contact between yeast and HeLa cells and incubated for 30 minutes on the benchtop. Unbound yeast were removed by five washes with PBS. Lysis buffer (10 mM EDTA + 0.1% Triton X-100 in PBS) was used to lyse mammalian cells, and yeast were diluted and plated on fresh YPD plates. Colony-Forming Units (CFUs) were then counted and compared to CFUs on input control plates, plated directly from yeast cultures at the same dilution.

Data were normalized between multiple experiments by division of each sample’s percent binding by the mean percent bound in the HA-Epa1p (3 rep) expressing strain of *S. cerevisiae*. We utilized the following equation to calculate display-normalized binding, which normalizes adherence based on Epa1p surface display metrics determined by flow cytometry.

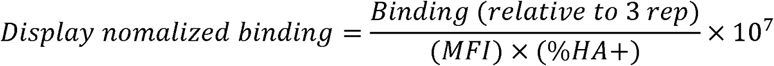

### Statistical Analysis

A Single Factor Analysis of Variance (ANOVA) was utilized to determine significant differences between experimental samples. Once ANOVA identified that there was variation between groups, one-tailed paired t-tests with Bonferroni Correction were run between all populations to determine which were the significantly different groups. The statistically significant α value was set at p<0.05.

## Results

### EPA1 codes for a repetitive and variable adhesin

To explore natural variation in *EPA1* repeat length, we amplified the repeat region from the genomic DNA of 24 clinical isolates and the BG14 reference strain by PCR. By agarose gel electrophoresis we observed four distinct band sizes, which corresponded to three (two isolates and BG14), four (seven isolates), five (14 isolates) repeats (Figure 1). Repeat length was confirmed via Sanger sequencing from a representative collection of clinical isolates and BG14 (Figure S2). One isolate (CG2) had a larger *EPA1* repeat region that could not be fully sequenced due to limits of Sanger sequencing read length; we calculated that this strain had ten tandem repeats in the *EPA1* repeat region by DNA migration (Figure 1, data not shown).

### HA-Epa1p surface display is modulated by repeat copy number

To study the effect of the *EPA1* repeat region on the characteristics of the protein, we designed a transgenic system to study Epa1p in the related yeast, *S. cerevisiae*. This orthogonal system was implemented as a means of normalizing expression of *EPA1* in a non-adherent background strain. HA-Epa1p variants with different repeat copy numbers were expressed in *S. cerevisiae* under the constitutive ADH promoter (Figure 1C). Surface display of HA-Epa1p with various Epa1p repeat copy numbers in *S. cerevisiae* was measured via flow cytometry (Figure 2). The percentage of cells stained positive for HA-Epa1p increases as Epa1p repeat copy number increases (Figure 2C). A one-way ANOVA was conducted between HA-Epa1p expressing strains, and there was found to be significant variation between groups (p<0.001), with the repeat region knockout (0 rep) having significantly fewer cells stained positive for HA (HA+) than all other repeat variants (p<0.01). When per cell density of HA-Epa1p on the cell surface was analyzed via MFI, the opposite trend was seen (Figure 2D). HA-Epa1p with the repeat region deleted had a significantly higher MFI than the repeat copy numbers (three, four, and five) seen in the natural population (p<0.0001).

### Heterologous EPA1 expression under constitutive expression decreases with increase repeat copy number

To study the underlying mechanisms which may control the variable Epa1p surface display we observed by flow cytometry, *EPA1* mRNA levels were measured by qPCR (Figure 3). In general, strains expressing *EPA1* under the ADH promoter expressed the transcript at a level 50-fold higher than under the native promoter in BG14 when grown in rich media (Figure 3A). Overall, increasing repeat copy number inversely correlated with *EPA1* mRNA transcript levels, with the five-repeat copy expressing at about 50% of the level of *EPA1* with the repeat region deleted (Figure 3B). While ANOVA found significant variation between the different copy number variants (p<0.01), the increase in expression of *EPA1* (0 rep) compared the other variants failed statistical significance. Rather, the only significant changes were found when comparing other variants to *EPA1* (10 rep) (p<0.05), which showed no expression in our system (Figure 3B).

### Repeat region protects HA-Epa1p from cell surface shedding

We then hypothesized that the differences surface display observed via flow cytometry could be explained by rapid degradation of Epa1p with fewer repeats. There is evidence for a protective role of protein glycosylation in related proteins (Karunanithi *et al.*, 2010, Meem & Cullen, 2012), and we wondered whether a key function of the repeat region might be to mediate this protection. To test this, we probed the growth media of HA-Epa1p expressing cells for the presence of HA-tagged fragments. As expected, when the media was analyzed via Western blot, there were two major HA+ protein species present, both migrating at apparent sizes smaller than the predicted cell wall associated protein size of ∼115 kDa (Figure 4A). These products ran at an apparent molecular weight of 70 kDa and 40 kDa, respectively and likely represent proteolytically cleaved products that are shed from the cell surface (Figure 4B). Qualitatively, as repeat copy number increases, there is a decrease in both fragments present in media.

### Deletion of the EPA1 repeat region impacts binding

HA-Epa1p was expressed in *S. cerevisiae* under the ADH promoter and bound to neuraminidase-treated HeLa cells to test the clinical relevance of differences in repeat copy number. The pADH negative control conferred no binding to epithelial cells while all three HA-Epa1p expressing strains tested showed binding at a similar level (Figure 5B). However, when epithelial cell binding was normalized to surface display metrics determined through flow cytometry, a different trend was seen (Figure 5C). HA-Epa1p with the repeat deleted performed significantly worse at the per protein level (p<0.05) when compared to HA-Epa1p with three or five repeats, which performed similarly to one another when display normalized binding was calculated (Figure 5C).

## Discussion

### Epa1p repeat copy number variation exists within the clinical population

Repetitive regions of DNA serve as important loci for population-wide strain to strain variability in yeast which leads to a variety of different phenotypes (Babokhov *et al.*, 2018), so we hypothesized that there would be variability in the copy number of the 120 bp *EPA1* minisatellite. Based on observations from other adhesins with similar repetitive domains, we hypothesized that binding would be improved by expanding repeat regions (Verstrepen *et al.*, 2005, Fidalgo *et al.*, 2006). Indeed, when we analyzed the region’s size in various clinical isolates by PCR, we found that there was a considerable amount of variation at this locus (Figure 1). We concluded that the majority of strains had an *EPA1* gene with between three and five repeats, with a single isolate (CG2) having ten repeats (Figure 1) and confirmed that the variation was the result of expansions of the tandem repeat region via sequencing (Figure S2).

In line with previous research in the field of repetitive domains in GPI-CWPs, there appears to be selection for Epa1p regions with expanded repeat regions. However, it was very interesting that there was only one strain (CG2) whose *EPA1* gene coded for a repeat region with greater than five repeats in Epa1p, suggesting that unchecked expansion of this repeat actually is detrimental for function.

### The tandem repeat region plays a role in regulating Epa1p display through multiple mechanisms

To test the functional differences of Epa1p with various repeat copy numbers, HA-tagged Epa1p with various repeat copy numbers were heterogeneously expressed in the related yeast, *S. cerevisiae*. Expressing HA-tagged Epa1p in a related species under a continuative promoter allowed us to uncouple our studies from the highly regulated gene expression of adhesin genes natively in *C. glabrata* (Timmermans *et al.*, 2018). This approach had additional advantages in that *S. cerevisiae* has no inherent adhesive properties toward mammalian cells (Lopez-Fuentes *et al.*, 2018) and shares a similar cell wall structure to *C. glabrata* (Cormack *et al.*, 1999, de Groot *et al.*, 2008).

To test how variation in the *EPA1* repetitive region affects phenotype, the cell surface display of HA-Epa1p was analyzed via flow cytometry (Figure 2). We tested HA-Epa1p variants with three, four, or five repeats, or with the repeat region deleted; the ten-repeat variant was omitted because we were unable to identify expression at the mRNA level (Figure 3). Significantly higher mean fluorescent intensity was observed for the cells expressing HA-Epa1p with the repeat region deleted than any of the naturally occurring repeat copy number variants (Figure 2D). This trend suggests there is decreased rate of protein surface display of Epa1p as a result of increased size of the repetitive region. However, HA-Epa1p with increased repeat copy number was displayed on a higher percentage of the total population (Figure 2C). Taken together, these findings suggest that variation of the Epa1p repeat region may exert control on surface display through multiple mechanisms.

To identify these mechanisms, we first measured the transcription of all of repeat variants (Figure 3). We observed that *EPA1* with increased repeat copy number expressed at lower levels than the gene with fewer copies of the tandem repeat. While not statistically significant, *EPA1* with the repeat region deleted appears to be more highly expressed, roughly 1.5-fold higher, than the native forms of the gene. Furthermore, no expression of *EPA1* (10 rep) was detected. This trend fits with previous findings that transcription of repetitive DNA is impaired as repeat copy number increases through a variety of mechanisms (Feng *et al.*, 2007, Biscotti *et al.*, 2015, Madireddy & Gerhardt, 2017). This trend may serve as an explanation as to why only one clinical isolate had a repeat copy number greater than five; without the expression of *EPA1*, pathogenicity of *C. glabrata* would be severely impaired.

While the difference in mRNA levels may explain the increased density of Epa1p on the surface of the cells when the repeat region is deleted, it does not explain why fewer of these cells are stained positively for HA-Epa1p. The repeat region is rich in glycosylated Ser/Thr residues which may contribute to stability within the yeast cell wall (Klis *et al.*, 2010, Meem & Cullen, 2012). To test this, we probed the cell culture media for differential loss of HA-Epa1p from the cell wall across repeat copy number variants (Figure 4). The total amount of HA-tagged product in the culture media decreases as repeat copy number increases, which suggests that indeed greater copy numbers stabilize Epa1p in the cell wall (Figure 4B). Neither of the fragments observed by Western Blot were the expected molecular weight of fully intact HA-Epa1p (∼115 kDa), supporting that HA-Epa1p is shed from the cell surface by cleavage by extracellular proteases. While our study did not directly assess the glycosylation status of the repeat region, we conclude that this is the most likely mechanism that drives increased stability in the cell wall as this process occurs for other GPI-CWPs (Karunanithi *et al.*, 2010, Meem & Cullen, 2012). We predict that as the repeat copy number, and therefore Ser-/Thr-associated glycosylation, increases, HA-Epa1p stability in the cell wall improves, resulting in less loss of HA-Epa1p via shedding.

### Binding capacity of Epa1p is decreased when the repeat region is deleted

Epa1p-mediated adherence to epithelial cells is an essential step in the establishing *C. glabrata* infection, and we looked to determine how repeat copy number would affect this process, given its complicated influence on Epa1p expression, display, and retention. When we tested the ability of our HA-Epa1p expressing *S. cerevisiae* to bind to sialic acid-depleted epithelial cells in an adherence assay based on those established by previous groups (Cormack *et al.*, 1999, Halliwell *et al.*, 2012, Diderrich *et al.*, 2015), we observed that transgenic expression of the adhesin confers the otherwise non-adherent yeast the ability to bind to epithelial cells in culture (Figure 5). Surprisingly, regardless of the Epa1p repeat copy number there was no trend between repeat copy number and adherence at the population level (Figure 5B). However, when we normalized the adherence to account for differences in surface display determined by flow cytometry, we found that HA-Epa1p (0 rep) confers less adherence than HA-Epa1p with naturally occurring repeat copy numbers (Figure 5C). This result supports data showing that decreased Epa1B domain size causes the decreased clearance of the cell wall by the ligand binding domain, suggests the repeat region may be required for cell wall clearance (Frieman *et al.*, 2002). However, with high density of the protein on the cell surface, the cell is able to overcome this obstacle as shown by equal adherence of non-normalized biding (Figure 5B). Even once surface display is normalized for, Epa1p variants with three and five repeats mediate adherence to epithelial cells with similar efficacy (Figure 5C).

Structural variation within the repeat region of Epa1p influences protein function by simultaneously influencing mRNA expression, protein trafficking, and protein turnover. In our heterologous system, these tradeoffs are functionally balanced such that only the extremes of repeat copy number (either no repeats or 10 repeats) demonstrate impaired Epa1p-mediated cell adhesion. While still speculation, differences in repeat copy number in the native *C. glabrata* likely have similar tradeoffs. Furthermore, it is also possible that repeat-length differences may mediate subtle differences which modulate cell adhesion, systemic migration, and cell wall integrity in the native system during infection, which we would be unable to detect in our system. Indeed, genomic analysis of the same *C. glabrata* infection before and after a course of antifungal treatment revealed lability in tandem repeat regions, perhaps especially so in the complement of cell wall proteins (Vale-Silva *et al.*, 2017). Thus, we predict that this variation may be a natural mechanism for tuning individual protein properties. As such, considering these mechanisms could prove crucial in treating infections of *C. glabrata* and understanding how certain infections achieve drug resistance.

## Acknowledgements

We thank Professor Brendan Cormack (Johns Hopkins University) and Dr. Yoav Golan (Tufts Medical School) for providing strains and clinical isolates which have been invaluable resources for this project. We also thank the Van Deventer lab for their assistance with Flow Cytometry. We thank Jessica Ziccarello, Jarett Mirecki, and Amanda Moises for their contributions to this project, and all other members of the Fuchs lab, past and present, for their helpful discussions and suggestions.

