## Supplementary material for "Variation in repeat copy number of the *Epithelial adhesin 1* tandem repeat region leads to variable protein display through multiple mechanisms": Table S1 and S2 and Figure S1 and S2

**Table S1.** *Oligonucleotides and SPHA cassette used in this study*

| Strain | Source |
|---|---|
| CG1 | blood |
| CG2 | blood |
| CG3 | blood |
| CG4 | blood |
| CG5 | bone |
| CG6 | dialysate |
| CG7 | blood |
| CG8 | RV lead |
| CG9 | perihepatic fluid |
| CG10 | aqueous fluid |
| CG11 | bladder tap |
| CG12 | abdominal fluid |
| CG13 | cornea |
| CG14 | aortic graft |
| CG15 | blood |
| CG16 | JP drain |
| CG17 | perihepatic fluid |
| CG18 | peritoneal fluid |
| CG19 | blood |
| CG20 | buttock tissue |
| CG21 | peritoneal dialysate |
| CG22 | pancreatic abscess |
| CG23 | hip synovial fluid |
| CG24 | blood |

**Table S2.** *Clinical sources of C. glabrata isolates*

| Name | Sequence (5'→ 3') |
|---|---|
| OCR014 | ATGATTTTAAATCCAGCTCTATTTTTG |
| OCR015 | TTAGGTCCCTATGTTCATC |
| OCR035 | GCCCGGATCCACTAGTTCTAGACCCTTAGGTCCCTATGTTCATC |
| OCR036 | ATATGATGTGCCAGACTACGCGCCCACATCTTCCAATGATATCAG |
| Epa1rep F | CACTGGAGCTCATCTGTCC |
| Epa1rep R | AACTAGCGTAACTGCTAGCC |
| EPA1 qPCR F | GATATTTCCAAAAATGGTAAGG |
| EPA1 qPCR R | GGGCTCAAAAACAGCTAAAG |
| ACT1 qPCR F | CCTTCAACGTTCCAGCCTTC |
| ACT1 qPCR R | CCGGCGTAAATTGGAACAAC |
| SPHA cassette | TACAATCAACTCCAAGCTGGCCGCTATGATTTTAAATCCAGCTCTATTT<br>TTGAATAAATGTGTATGCATTTACACTACTCTCATTTTGCTCCTATTAAC<br>GAACGGTGGTTATGCTACATCTTCCTACCCCTATGACGTTCCCGACTA<br>CGCTTATCCTTACGACGTACCTGATTACGCCTACCCATATGATGTGCC<br>AGACTACGCGCCCGGGTCTAGAACTAGTGGATCCGGGCTGCAGGAAT<br>TCGATATCAAGC |

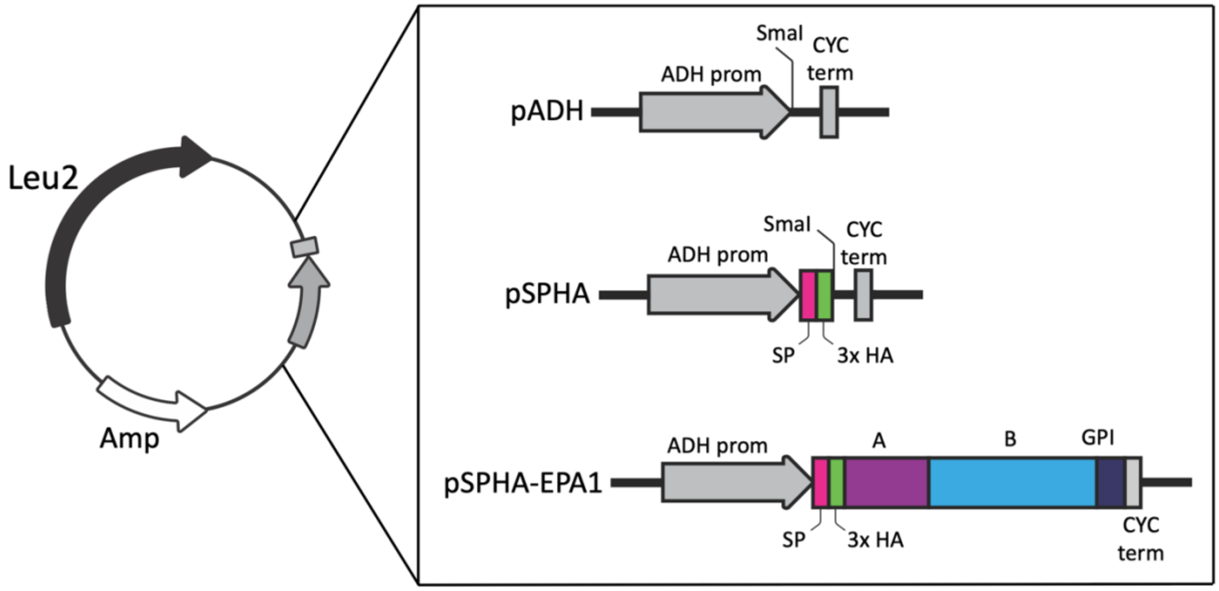


**Figure S1.** *Plasmid construction*. All plasmids were constructed from the pADH (Addgene 87374) backbone which codes for leucine selection [Leu2] in yeast and ampicillin resistance (Amp) in *E. coli*. pADH also contains the ADH promotor and CYC terminator for constitutive expression of a gene of interest in yeast. To construct the pSPHA surface expression vector, the *EPA1* cell surface trafficking signal peptide sequence (SP), a 3xHA tag, and multiple cloning site were cloned into pADH at the SmaI restriction site. The segment of the *EPA1* gene which codes for the A and B domains and GPI consensus sequence were then cloned into the SmaI site of pSPHA to form pSPHA-EPA1, which enables yeast to express and HA-tagged Epa1p on the surface of the cell.

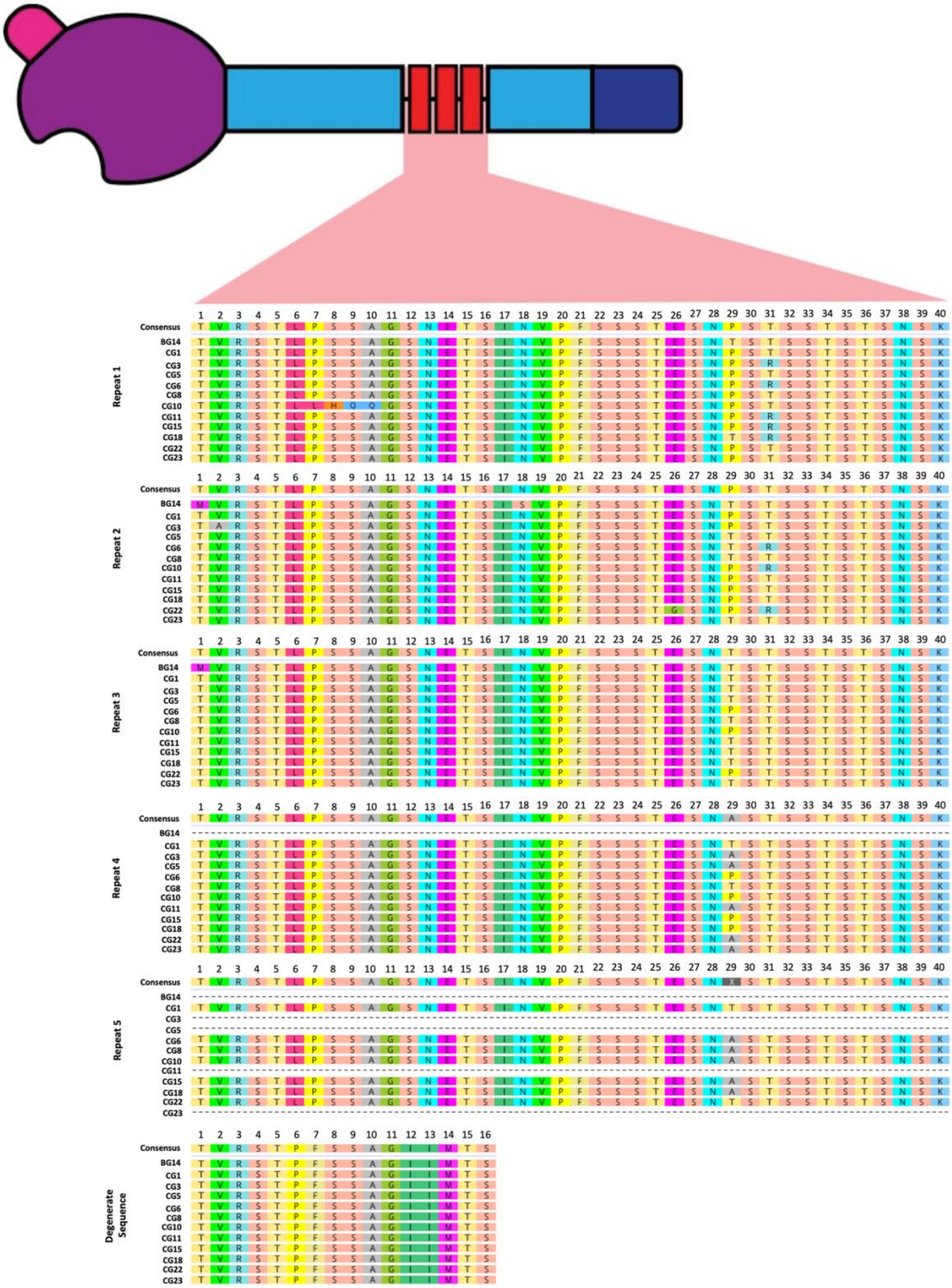


**Figure S2.** *Protein alignment of Epa1p repeat sequences from representative sample of C. glabrata isolates*. *EPA1* was isolated from the gDNA of select *C. glabrata* isolates (CG1-24) and the BG14 reference strain and the repetitive region was sequenced.
